# GIGEM (Group Isolation Gauge Effect Metrics), A Software Suite for Analyzing Social Isolation-Induced Sleep Loss and Multi-Batch Experiments in *Drosophila*

**DOI:** 10.64898/2025.12.18.695326

**Authors:** Esther I. Doria, Feda Hammood, Jingjing Yan, Wanhe Li

## Abstract

We introduce Group Isolation Gauge Effect Metrics (GIGEM), a high-throughput software suite for *Drosophila* sleep analysis designed to manage large population datasets from multi-batch experiments. In this study, we characterized social isolation-induced sleep phenotypes across 38 Sleep Inbred Panel (SIP) lines after both acute and chronic social isolation. Beyond this specific study, GIGEM is broadly suited for diverse *Drosophila* sleep research involving large-scale and multi-batch datasets. GIGEM enabled an extensive analysis of sleep parameters and allowed systematic identification of sleep changes in response to social isolation. Following acute social isolation, SIP line animals showed heterogeneous responses, including sleep loss, sleep gain, and no change in sleep. In contrast, chronic social isolation led to significant sleep loss in nearly all lines, though to varying degrees. While most SIP lines were sensitive to chronic social isolation, a select few were resistant to it. These results were obtained across 38 SIP lines and wild-type controls (totaling 5670 animals). Collectively, our results demonstrate that chronic social isolation-induced sleep loss is a robust yet variable trait and establish GIGEM as a powerful, generalizable tool for *Drosophila* sleep phenotyping in large-scale, multi-batch studies.

## INTRODUCTION

Health problems arising from chronic social isolation are a major public health concern. The adverse health effects of social isolation are thought to have a behavioral basis, particularly involving sleep deficits, which are associated with vascular disease, metabolic syndrome, diminished immunity, and depressive symptoms [1–4]. Understanding how social isolation affects sleep behavior is, therefore, critical. Recent studies have used the genetic model organism, *Drosophila melanogaster*, to study social isolation-induced sleep loss. In laboratory wild-type *Drosophila* strains, social isolation induces sleep loss in a duration-dependent manner. Chronic social isolation (5-7 days) leads to a significant sleep loss, while acute social isolation (1-3 days) does not [5–7]. This robust, reproducible phenotype serves as an ideal platform for exploring the genetic basis of behavioral changes induced by social isolation, leveraging the extensive genetic resources and high-throughput capacity of the *Drosophila* model.

Large-scale multi-batch studies are essential for investigating genetic effects of social isolation-induced changes in sleep, but analyzing such datasets presents major challenges: experimental batch effects, complex multi-parameter outputs, and the need for consistent cross-strain and/or experiment comparisons. To address these issues, we developed the Group Isolation Gauge Effect Metrics (GIGEM), an R-based software for high-throughput analysis of *Drosophila* sleep behavior. GIGEM includes a basic batch analysis module for processing individual experimental sets and a multi-batch analysis module for integrating results across experiments.

To demonstrate GIGEM’s capabilities, we analyzed social isolation-induced sleep phenotypes in 38 lines from the Sleep Inbred Panel (SIP), a collection of 39 inbred *Drosophila melanogaster* strains with naturally occurring polymorphisms and stable long- or short-sleeping traits [8, 9]. We hypothesized that variation in sleep duration and architecture would lead to differences in isolation-induced sleep loss among SIP lines. The 38 SIP lines were divided into batches and tested alongside a *Canton S* (CS) control strain under acute (2-days) and chronic (5-days) group and isolation conditions in a large, multi-batch experiment conducted over three months. GIGEM enabled extensive quantification of diverse sleep parameters, systematically identifying strain-specific responses and revealing chronic social isolation-induced sleep loss as a widespread but variable trait. Its multi-batch module integrated and adjusted results across experiments, allowing comprehensive comparisons among SIP lines to uncover trends and correlations in sleep phenotypes. This method thus provides a robust and scalable solution for large-scale sleep phenotyping in *Drosophila* and can be applied to diverse experimental paradigms.

## MATERIAL AND METHODS

### *Drosophila* Strains and Culture

*Drosophila melanogaster* stocks were raised on *Drosophila* culture media (cornmeal/yeast/molasses/agar) at 21.5°C under 12-hour light/12-hour dark (LD) cycles. The Sleep Inbred Panel (SIP) lines were obtained from the Bloomington Drosophila Stock Center. 38 of the 39 SIP lines were tested, as 1 SIP line had difficulty surviving in our laboratory environment. The wild-type isogenic strain *Canton S* (CS) was used as a control.

### Group Enrichment/Social Isolation

Newly eclosed fruit flies were collected and kept in standard fly food bottles at a density of ∼200 flies per bottle for 3–5 days to acquire social experience. Mating was allowed to happen freely during this period. Male flies were then sorted into standard fly food vials: 1 fly per vial for social isolation (Iso) and 25 flies per vial for group enrichment (Grp). The fly sorting day was considered Day0. On Day2 or Day5, group-treated and isolated flies were used for sleep measurements using the Drosophila Activity Monitors (DAM; Trikenetics Inc., Princeton, MA) (Figure 1A).

**Figure 1.**
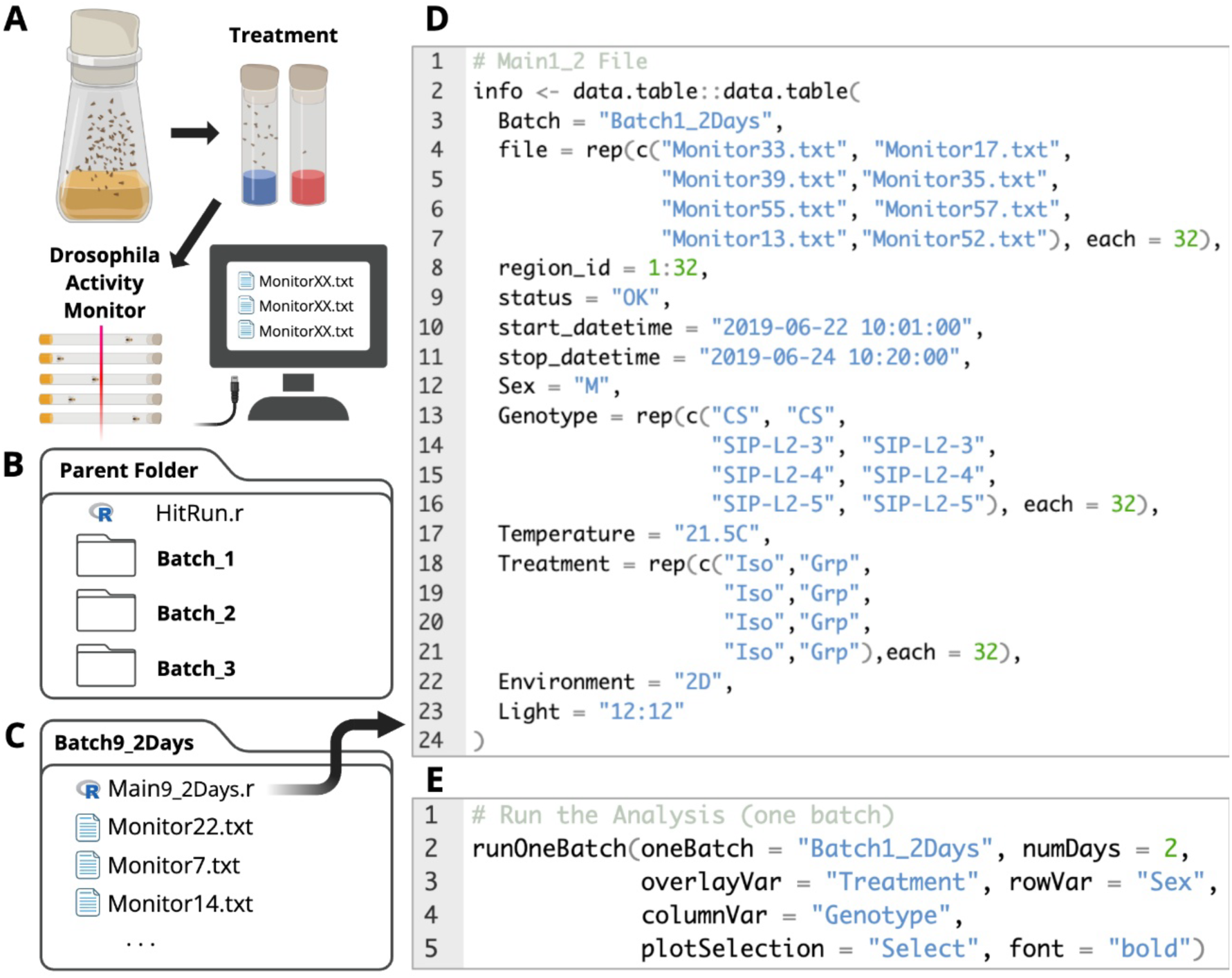
Experimental workflow. **(A)** Flies with social experience were collected and put through group (25 flies in a vial) or isolation (a single fly in a vial) treatments for 2 or 5 days and then monitored using Drosophila Activity Monitors (DAM). **(B)** The parent folder of GIGEM contains a HitRun.r file and batch folders. **(C)** Each batch folder contains the monitor files collected for the experimental batch and a Main.r file containing the experimental design. **(D)** The Main.r file describes the title of the batch, monitor information and experimental factors.

### Sleep Measurement and Analysis

Flies with group or isolation experience were loaded into glass tubes containing fly food and assayed at 21.5°C under 12-hour light/12-hour dark (LD) cycles. Activity counts were collected every minute for 2 LD cycles following the loading day, using DAM. Sleep analysis was conducted using the GIGEM software.

### GIGEM – Overall Features

The Drosophila Activity Monitor (DAM) is a high-throughput behavior platform for recording locomotor activity. However, analyzing sleep behavior from the raw data collected by DAM is challenging, and developing software to aid the analysis remains an ongoing community effort [10–16]. Our software, Group Isolation Gauge Effect Metrics (GIGEM), specializes in analyzing multiple batches of sleep experiments in parallel and exploring the data using fitted linear models and correlations. This capability is beneficial for analyzing social isolation-induced sleep changes in the SIP lines for several reasons. First, we had to separate the 38 SIP strains into multiple experimental batches, with each batch containing on average 338 isolated and group-treated animals. For each batch, the duration of isolation/group treatment was either 2 or 5 days. A total of 18 batches of data were analyzed in parallel, making multi-batch analysis a necessity. Second, a control wild-type strain must be included for each batch as behavior experiments often exhibit variation over time. The same control strain—in this case, CS—served as a uniform reference to compare SIP behavioral phenotypes. This consistent reference enabled us to fit a linear model to control for batch effects and to predict the treatment effect of social isolation for each strain. Third, assessing correlations between sleep traits and clustering data across all SIP strains can clarify the relationships among the SIP strains’ sleep behavior phenotypes.

Although GIGEM was tailored to analyze group/isolation treatment, it is also widely applicable to behavioral data from other types of treatments. GIGEM includes five functions: two for analyzing individual experimental batches and three for analyzing data across multiple experimental batches.

### GIGEM – Analysis Setup

GIGEM requires a specific computer file structure, with each batch in a separate folder nested inside a parent folder (Figure 1B). Each batch folder contains the monitor files collected by the DAM and a “Main.r” R file describing the experimental design (Figure 1C, D). The parent folder contains the “HitRun.r” R file and serves as the working directory. It is also where all multi-batch analysis outputs are saved. An example setup using the SIP-line data can be found at https://wanheli2022.wixsite.com/wanheli/gigem.

### GIGEM – Basic Functions

There are two basic functions in GIGEM to analyze sleep using raw data from DAM: runOneBatch and runAllBatches, which are built based on the R/rethomics [12], and R/dplyr [17], R/ggplot2 [18], R/ggbeeswarm [19], and R/ggprism [20] packages.

The runOneBatch function executes a single, explicitly named batch, while the runAllBatches function iterates through all batch folders within the parent directory and produces CSV files and plots containing sleep analysis results specific to each batch. When using the basic functions, a variable, “numbDays,” allows users to specify the number of days of DAM data used for analyses. Somnograms for all individuals within a batch are automatically generated for each monitor, both before and after removing dead flies from the dataset [12]. In addition, three CSV files are automatically generated. The first file, “loadinginfo.csv,” contains the list of animals and their associated experimental conditions for each experimental factor, as specified in the “Main.r” experimental design file (start/stop dates, genotype, treatment, etc.). The second file, “stat.csv,” provides summary statistics (including mean, standard deviation, standard error, and confidence interval) for each sleep parameter measured across the population. The sleep parameters of interest are: Sleep_Time_All (total sleep time during day and night), Sleep_Time_L (sleep time during the day/light phase), Sleep_Time_D (sleep time during the night/dark phase), n_Bouts_L (number of sleep bouts during the day/light phase), mean_Bout_Length_L (average bout length during the day/light phase), n_Bouts_D (number of sleep bouts during the day/light phase), and mean_Bout_Length_D (average bout length during the day/light phase). The third file, “summary.csv,” summarizes the sleep parameters listed above, along with a few additional sleep parameters: sleep fraction, sleep latency, first bout length, and latency to the longest bout for each individual animal. For each parameter, we report the mean across all monitored days and the corresponding values for each individual day (Day 1 through *numbDays*).”

For plotting the results, the variables “overlayVar,” “columnVar,” and “rowVar” are used to color and organize the experimental factors within the plots. The “columnVar” and “rowVar” divide plots into columns and rows, respectively, based on differing conditions within the specified experimental factors. In addition, the variable “plotSelection” can be set to “All”, “None,” or “Select” to determine which plots to generate. If “Select” is chosen, the program prompts the user with yes-or-no questions for a series of possible plots:

1. *Activity and Sleep Actograms*: Displays individual animals’ activity and sleep (inactivity) patterns over time.
2. *Population Sleep Profiles*: Shows the average percentage of time spent sleeping in consecutive 30-min time bins of a population of animals for a specified number of days, as well as a 24-hour “wrapped” average across those days (Supplementary Figure 1a).
3. *Overlaid Sleep Profiles*: Similar to *Population Sleep Profiles*, but with multiple profiles superimposed and color-labeled based on the experimental factor “overlayVar” is set to.
4. *Overlaid Sleep Bout Profiles*: Displays each population’s average sleep bout length averaged across analysis days and presented in a 24-hour wrapped format, with color-labeled profiles superimposed by condition in the experimental factor that “overlayVar” is set to.
5. *Quantifications of Sleep Parameters*: Generates three bar plots displaying the average amount (in minutes) of daytime (light) sleep, nighttime (dark) sleep, and total sleep; and four violin plots displaying the average length of sleep bouts and the average number of sleep bouts during both light and dark phases.
6. *Combined Plots (Within Batches)*: Groups the “wrapped” *Overlaid Sleep Profiles* and *Quantifications of Sleep Parameters* by each unique combination of experimental factors, continuing to use color labels from “overlayVar.”
7. *Combined Plots (Across Batches)*: Similar to *Combined Plots (Within Batches)*, but uses data across all batches. It outputs plots into the parent folder. If there are replicates of the same combinations of conditions in separate batches, it groups data from matching experimental conditions across batches. This plot is only available when running runAllBatches and is automatically set to “no” when running runOneBatch.

### GIGEM – Multi-Batch Functions

There are three functions specializing in analyzing sleep data from multiple batches as a whole: rankedDisplay, corMatrix, and corScatter.

The rankedDisplay function (Figure 2) generates a barplot of population averages across all batches for a specified experimental factor. The bars represent total, daytime, or nighttime sleep and are computed in one of the following three ways: (1) as the raw sleep amount in minutes, (2) as the difference between two specified conditions (𝑐𝑜𝑛𝑑𝑖𝑡𝑖𝑜𝑛1 − 𝑐𝑜𝑛𝑑𝑖𝑡𝑖𝑜𝑛2), or (3) as the percentage change between two conditions, calculated as (𝑐𝑜𝑛𝑑𝑖𝑡𝑖𝑜𝑛1 − 𝑐𝑜𝑛𝑑𝑖𝑡𝑖𝑜𝑛2) ÷ (𝑐𝑜𝑛𝑑𝑖𝑡𝑖𝑜𝑛2). If no conditions are specified, the function defaults to the first option. When conditions are specified, setting the variable “method” to “Diff” or “Perc.Change” leads to the second or third option, respectively. Optionally, setting “fitted = TRUE” causes the program to fit a linear model and use the linear model-predicted values instead of the actual raw values. This “fitted” variable can be used in conjunction with the percentage change method. If needed, the multiple conditions from experimental factors can be reduced to a single condition by subsetting, thereby eliminating that experimental factor from affecting the linear model by limiting the linear model’s scope of analysis. The linear model uses parameters defined by an equation that the user sets with the “formula” variable. For example, in this paper, the linear model for rankedDisplay, corMatrix, and corScatter was set as a function of “Genotype * Treatment + Batch” for 2D and 5D independently. The linear model-predicted values for each genotype were then used to calculate the sleep change using the “Diff” method, with condition 1 and condition 2 set to social isolation and group treatments, respectively.

**Figure 2.**
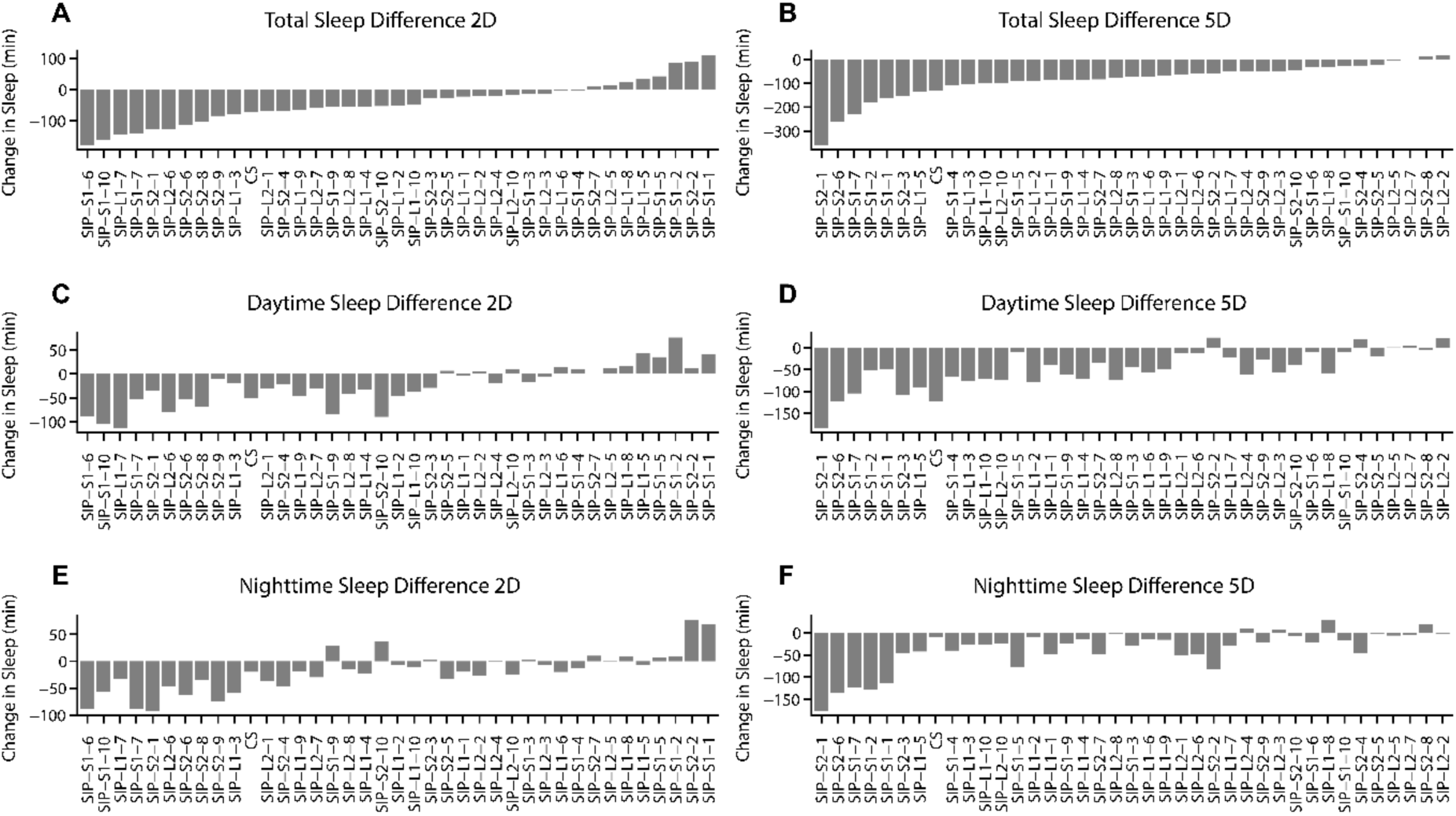
Social isolation-induced sleep changes in the SIP lines. The data was filtered for outliers, fit to a linear model, and adjusted for batch effect to generate predicted values of sleep parameters for each strain. Each bar represents the change in sleep relative to the grouped condition for each strain, calculated as (*i* − *g*), where *i* is the isolated average and *g* is the group-housed average. (**A and B**) Total Sleep Change. **(C and D)** Daytime Sleep Change. **(E and F)** Nighttime Sleep Change. The ranking order is determined based on the social isolation-induced change in total sleep.

The corMatrix function generates a correlation matrix (Figure 3) of sleep parameters or traits based on whether the condition1 and condition2 variables have been set, with *p*-values adjusted for the false-discovery rate using the Benjamini-Hochberg method. Correlation matrices can also be generated from raw values or predicted values after a linear model has been applied.

**Figure 3.**
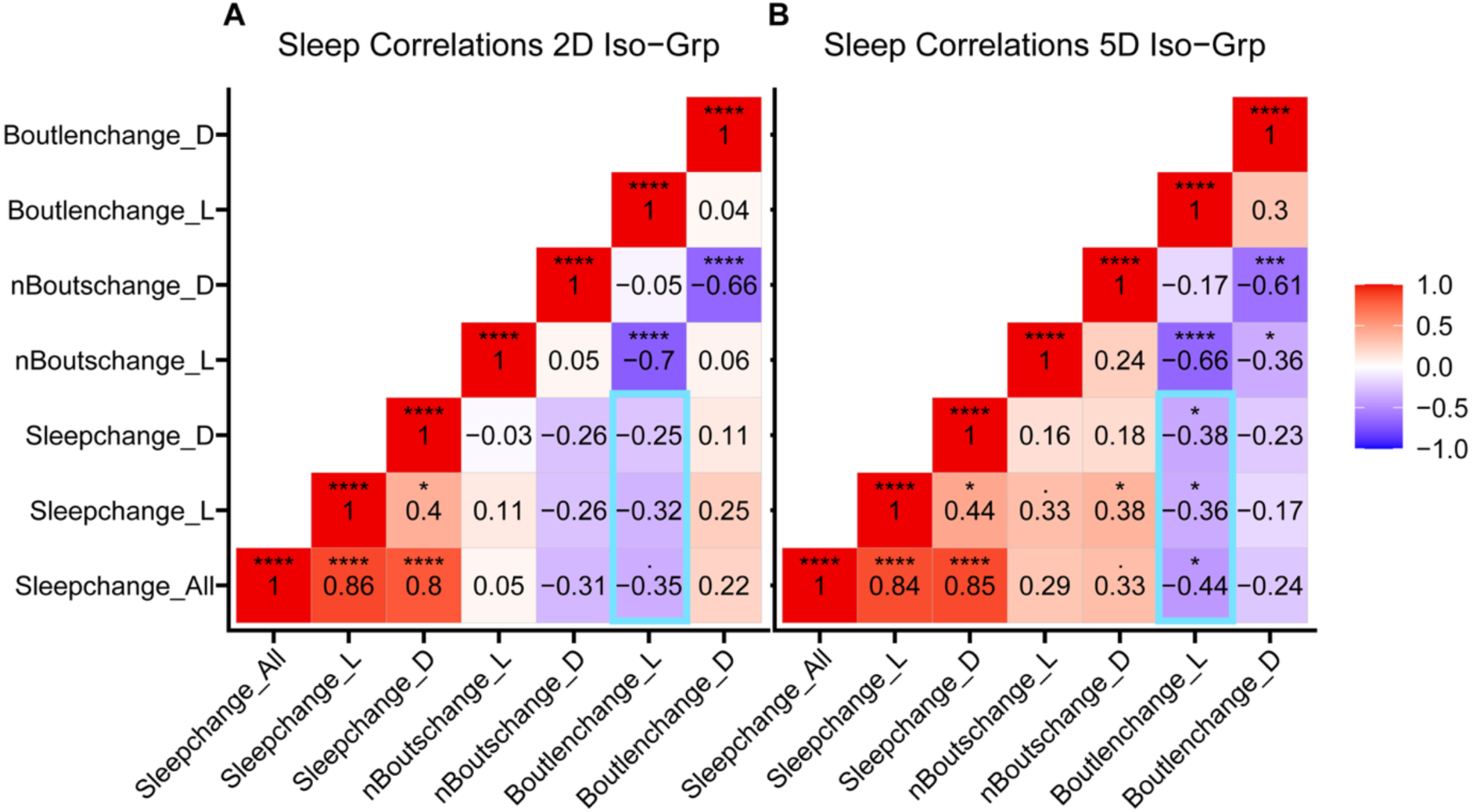
Correlation matrix of change in sleep parameters. The data was filtered for outliers, fit to a linear model, and adjusted for batch effect to generate predicted sleep values of sleep parameters for each strain. Each point represents the population average change in a given sleep parameter for a single strain, calculated as (*i* − *g*), where *i* is the isolated average and *g* is the group-housed average for that parameter. The dataset includes 38 Sleep Inbred Panel (SIP) lines and *Canton S* (CS), and is divided by 2-day **(A)** and 5-day **(B)** social isolation treatments. Pearson correlation coefficients (*r*) are displayed and heat-mapped within each matrix cell. Emerged correlations are highlighted. Statistical significance was assessed by two-sided unpaired *t*-tests and adjusted for false discovery rate using the Benjamini-Hochberg method, with thresholds of *\*P<0.05, **P<0.01, ***P<0.001, ****P<0.0001*.

The corScatter function (Supplementary Figure 2) provides a visualization of the data displayed from corMatrix by generating scatter plots of sleep parameters or traits. This facilitates visual data exploration, which may be hidden beneath the correlation matrix. As with rankedDisplay and corMatrix, corScatter can be produced using either raw values or linear-model-predicted values.

### GIGEM – Statistics and Reproducibility

Two-tailed unpaired *t*-tests were used to compare group and isolation treatments for all seven basic sleep parameters: Sleep_Time_All, Sleep_Time_L, Sleep_Time_D, n_Bouts_L, mean_Bout_Length_L, n_Bouts_D, and mean_Bout_Length_D. For multi-batch plots, outliers were filtered above 3 standard deviations from the strain-treatment population mean. The Pearson product-moment correlation test [21] was used for the correlation matrix, and p-values were adjusted for false-discovery rate using the Benjamini-Hochberg method in the correlation matrix (Figure 3). Statistical significance thresholds for all statistical tests were set to \**P*<0.05, \*\**P*<0.01, \*\*\**P*<0.001, and \*\*\*\**P*<0.0001.

## RESULTS

38 of the 39 total SIP lines were viable in the laboratory and used to demonstrate the usability of GIGEM. The SIP lines were divided into SIP-L1, SIP-L2, SIP-S1, and SIP-S2 lines, based on the genetic similarity as determined by the construction process of the SIP lines [8, 9]. SIP-L_ refers to long-sleep phenotypes while SIP-S_ refers to short-sleep phenotypes. The entire experiment contains 18 batches of data. We used runOneBatch, runAllBatch, rankedDisplay, corMatrix, and corScatter functions to analyze the experiment. A total of 5,680 flies were included in the analyses.

### Single Batch Analysis

Here, we demonstrated how to use GIGEM to conduct single batch analysis by measuring sleep of CS, SIP-L2-1, SIP-L2-2, SIP-S2-1, SIP-S2-8, and SIP-S2-10 lines after group/social isolation for 2 or 5 days. These experiments were designated as Batch9_2Days and Batch9_5Days in our dataset and included a total of 777 animals. We analyzed these two batches using the runOneBatch function and the results were summarized in Supplementary Figure 1.

Consistent with previous findings [5], wild-type *Canton S* (CS) flies exhibited significant sleep loss during daytime after chronic social isolation (5 days), which also led to a reduction in total sleep. Acute social isolation (2 days) had no significant effect. Although the sleep loss phenotype is distributed during daytime and is isolation duration-dependent in wild-type flies, social isolation-induced sleep loss appeared to be a variable trait among SIP lines. For example, SIP-L2-1 flies exhibited nighttime, rather than daytime, sleep loss after both acute and chronic social isolation. However, sleep fragmentation occurred in SIP-L2-1 after chronic social isolation, indicated by an increase in the number of sleep bouts and a decrease in mean bout length during the night phase. Sleep in SIP-L2-2 animals appeared unaffected by either 2 days or 5 days of isolation. SIP-S2-1 animals exhibited severe chronic social isolation-induced sleep loss during total, light, and dark phases. SIP-S2-8 and SIP-S2-10 exhibited a reversed pattern of social isolation duration-dependent sleep loss compared to CS. Acute social isolation-induced daytime sleep loss, but chronic social isolation did not cause any change in sleep (Figure 2 and Supplementary Figure 1). These results suggest social isolation-induced change in sleep is a prevalent but variable trait.

### Multi-Batch Analysis

We used the runAllBatches function to analyze the multi-batch dataset, which included a total of 5,680 animals tested for sleep after group treatment or social isolation for 2 or 5 days. The analysis results for sleep profiles and all sleep parameters were summarized in Supplementary Figure 3–6. We next demonstrated how to use GIGEM to investigate all SIP strains together by applying the rankedDisplay, corMatrix, and corScatter functions.

Using the rankedDisplay function, social isolation-induced sleep loss is displayed in a ranked way across the 38 SIP lines (Figure 2). A linear model was applied to take batch effect into consideration, and the linear model-predicted values were used for the ranking. The range of sleep change (𝐼𝑠𝑜𝑙𝑎𝑡𝑖𝑜𝑛 𝑠𝑙𝑒𝑒𝑝 − 𝐺𝑟𝑜𝑢𝑝 𝑠𝑙𝑒𝑒𝑝) after 5 days of social isolation varied from -359 minutes to +28 minutes. The range of sleep change after 2 days (𝐼𝑠𝑜𝑙𝑎𝑡𝑖𝑜𝑛 𝑠𝑙𝑒𝑒𝑝 − 𝐺𝑟𝑜𝑢𝑝 𝑠𝑙𝑒𝑒𝑝) of social isolation was smaller than that after 5 days of social isolation. While 35 of 38 SIP lines exhibited chronic social isolation-induced sleep loss, 30 of 38 SIP lines exhibited acute social isolation-induced sleep loss. Neither social isolation-induced sleep loss distributed during daytime and/or nighttime appeared to be strain-specific. These results further demonstrated that social isolation-induced sleep loss is a prevalent but variable trait, and the multi-batch analysis in GIGEM can facilitate exploring its variability.

Using the corMatrix and corScatter functions, we calculated isolation-induced changes in seven sleep parameters for each strain and examined how these changes correlated across all strains (Figure 3 and Supplementary Figure 2). These calculated changes included: change in total sleep time (Sleepchange_All), change in daytime/nighttime sleep (Sleepchange_L and Sleepchange_D), change in the number of daytime/nighttime sleep bouts (nBoutschange_L and nBoutschange_D), change in daytime/nighttime average sleep bout length (Boutlenchage_L and Boutlenchage_L). For each sleep parameter, the change was calculated as: the predicted values of the sleep parameter of the group-treated animal – the predicted values of the sleep parameter of isolated animal.

Most correlations between isolation-induced changes appeared consistent, with similar *p*-values after two or five days of isolation. Change in total sleep time was strongly correlated with both change in daytime and nighttime sleep. As expected, the change in the number of sleep bouts was always negatively correlated with the change in average sleep bout length, during both daytime and nighttime (Figure 3).

Importantly, several new correlations emerged after 5 days of social isolation: change in daytime bout length exhibited negative correlations with change in total/nighttime sleep after 5 days of social isolation, but not after 2 days of social isolation (Figure 3B). These emerged correlations suggested that chronic social isolation altered the circadian phase distribution of sleep behavior and demonstrated that it is important to explore the multi-batch data using correlation analysis.

## DISCUSSION

Sleep research produces extensive time-series datasets, and the increasing prevalence of high-throughput experimental designs places substantial demands on data analysis. To meet these challenges, GIGEM provides streamlined, multi-batch analytical tools that facilitate large-scale experiments encompassing numerous treatment groups over extended durations. In this study, we studied social isolation-induced sleep loss using the Sleep Inbred Panel (SIP) strains to demonstrate both the single-batch and multi-batch capabilities of GIGEM. The single-batch functionality helped us analyzed sleep of 5,680 animals efficiently, and the multi-batch functionality allowed adjustment of batch effect and data exploration. We concluded that social-isolation-induced sleep loss is a prevalent but variable trait. We observed emerging correlations between change in daytime bout length and change in sleep, as well as between change in nighttime sleep bouts number and change in daytime sleep. These correlations warrant further validation using additional strains beyond the SIP lines to assess their generalizability and explore the genetic basis of social isolation-induced behavioral changes.

GIGEM enables efficient analysis of large-scale sleep datasets and exploration across multiple strains and conditions. While currently configured for social isolation experiments, it remains flexible for other high-throughput, multi-treatment designs. Importantly, GIGEM allows correction of batch effects and preparation of sleep parameters for correlation analysis, which are essential for downstream studies, such as genome-wide association studies using variable sleep traits, to reveal biological mechanisms underlying sleep regulation.

## Acknowledgements

We thank Fei Zhang, Daniel Nguyen, Daniel Doria, and Mateo Langston-Smith for advice on programming and statistical methods. We thank Zikun Wang, Laurel Eckhardt, and Michael Young for their help with experiment design and sharing equipment. We thank Xin Chen, Doris Migliaccio, Sydney Christensen, and Min Feng for comments on the manuscripts. We thank members of the Li laboratory for technical support. We thank the Bloomington Drosophila Stock Center for fly stocks. This work was supported by the Cancer Prevention and Research Institute of Texas [RR220021] and the National Institute of General Medical Science [GM150832] to W. L.. F. H. was supported by the Yalow Scholars Program and the McNulty Scholars Program at Hunter College.

## Author contributions

Conceptualization: W.L., F.H., and E.I.D.; data curation: E.I.D., F.H. and W.L.; formal analysis: E.I.D.; funding acquisition: W.L.; investigation: E.I.D., F.H. and W.L.; methodology: E.I.D., F.H., and W.L.; software: E.I.D. and W.L.; validation: J.Y.; supervision: W.L.; visualization: E.I.D.; writing-draft: E.I.D. and W.L.; writing-editing: E.I.D. and W.L.. All authors have read and agreed to the published version of the manuscript.

## Data Availability

The GIGEM package developed, and all data used in this study can be downloaded through https://wanheli2022.wixsite.com/wanheli/gigem. Analyzed data with sleep parameters from all batches are available in Supplementary Figures 3-6.

## Supplementary Materials

**Supplementary Figure 1.**
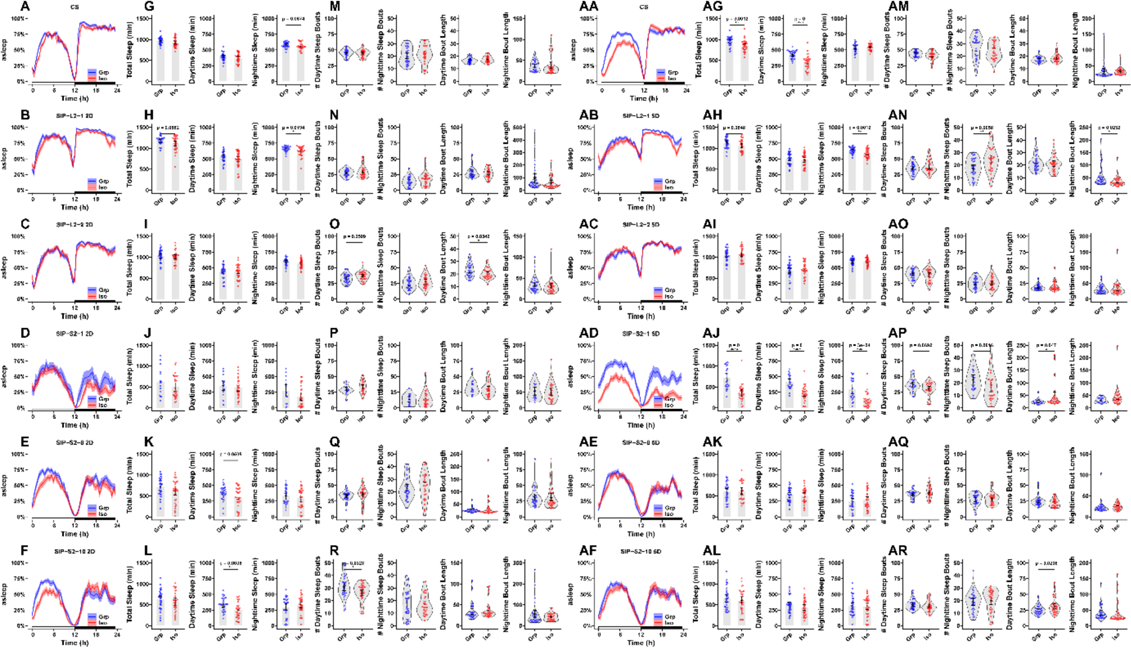
Sleep profiles and sleep parameters for a single experimental batch. **(A–F; AA–AF)** Population sleep profiles show the percentage of animals asleep in 30-minute intervals, averaged across 2 LD cycles after isolation treatment of 2 days (A-F) or 5 days **(AA–AF)**. Solid lines represent the mean; shaded areas indicate ±SEM. Each data point represents an individual animal’s average for the indicated sleep parameter, including: total sleep (ZT0–24) daytime sleep (ZT0–12), and nighttime sleep (ZT12–24) **(G–L; AG–AL)**, as well as the number of daytime sleep bouts, the number of nighttime sleep bouts, average daytime bout length, and average nighttime bout length **(M–R; AM–AR)**. Social isolation-induced sleep changes after 2 days of social isolation are shown on the left. Social isolation-induced sleep changes after 5 days of social isolation are shown on the right. Statistical significance was assessed by two-sided unpaired *t*-tests (*\*P<0.05, **P<0.01, ***P<0.001, ****P<0.0001*; *n* = 15–32 flies.

**Supplementary Figure 2.**
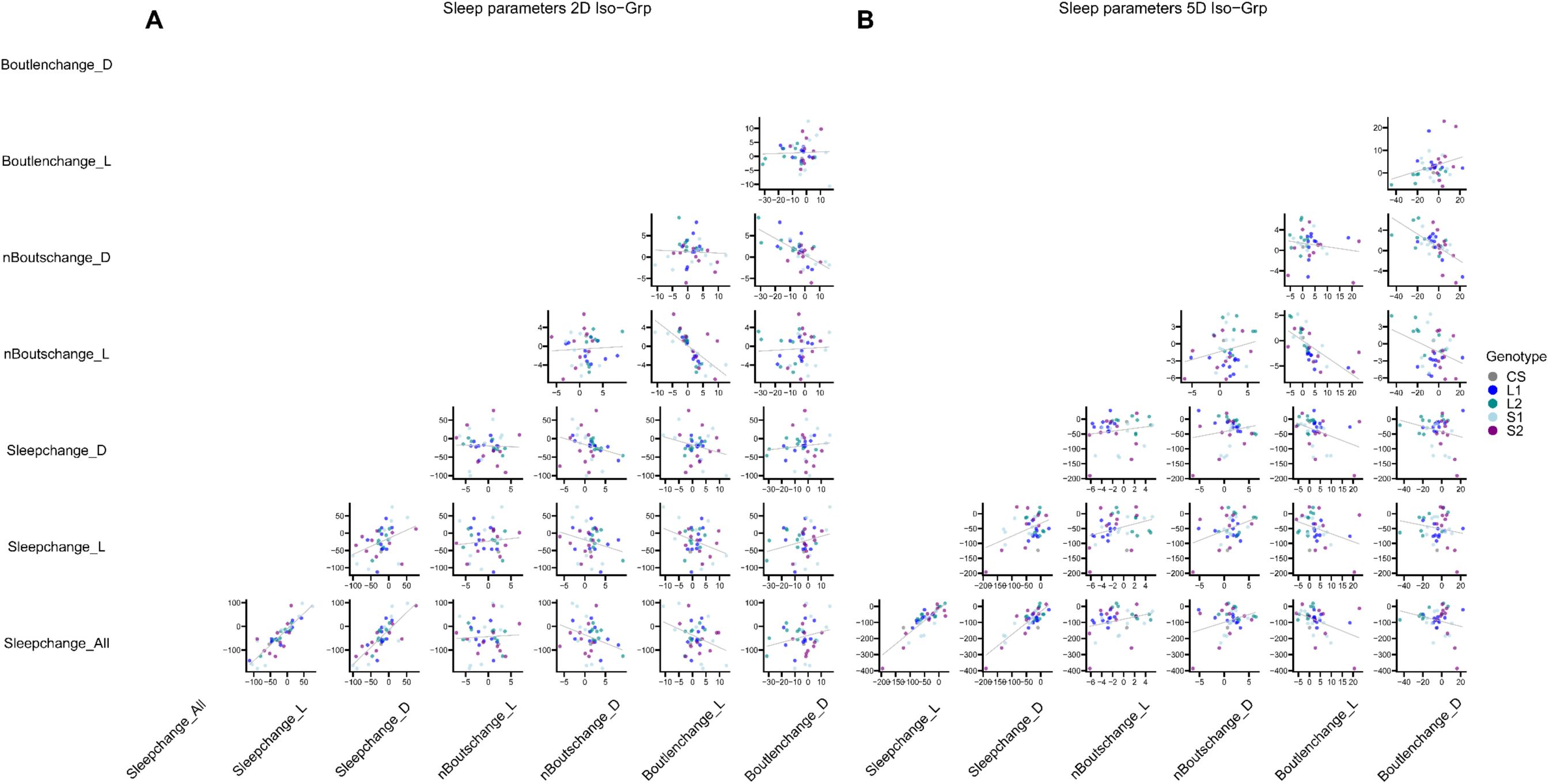
Scatterplot visualization of correlation matrix data (Figure 3). The data was filtered for outliers, fit to a linear model, and adjusted for batch effect to generate predicted sleep values for each strain. Panels **A** and **B** display a matrix of scatterplots comparing the difference between isolated and group-housed averages (*i* − g) for each parameter by strain. Each point represents the strain-level mean for 38 Sleep Inbred Panel (SIP) lines and *Canton S* (CS). Linear regression lines are overlaid to indicate the direction and center-line of the relationships between traits. Colors represent genetic background. Data is divided by social isolation duration for 2 days (A) or 5 days (B). These plots visualize the same data shown in the correlation matrices of Figure 3, with scatterplot positions corresponding directly to the correlation matrix’s boxed *r*-value position. *n* = 39 strain

**Supplementary Figure 3.**
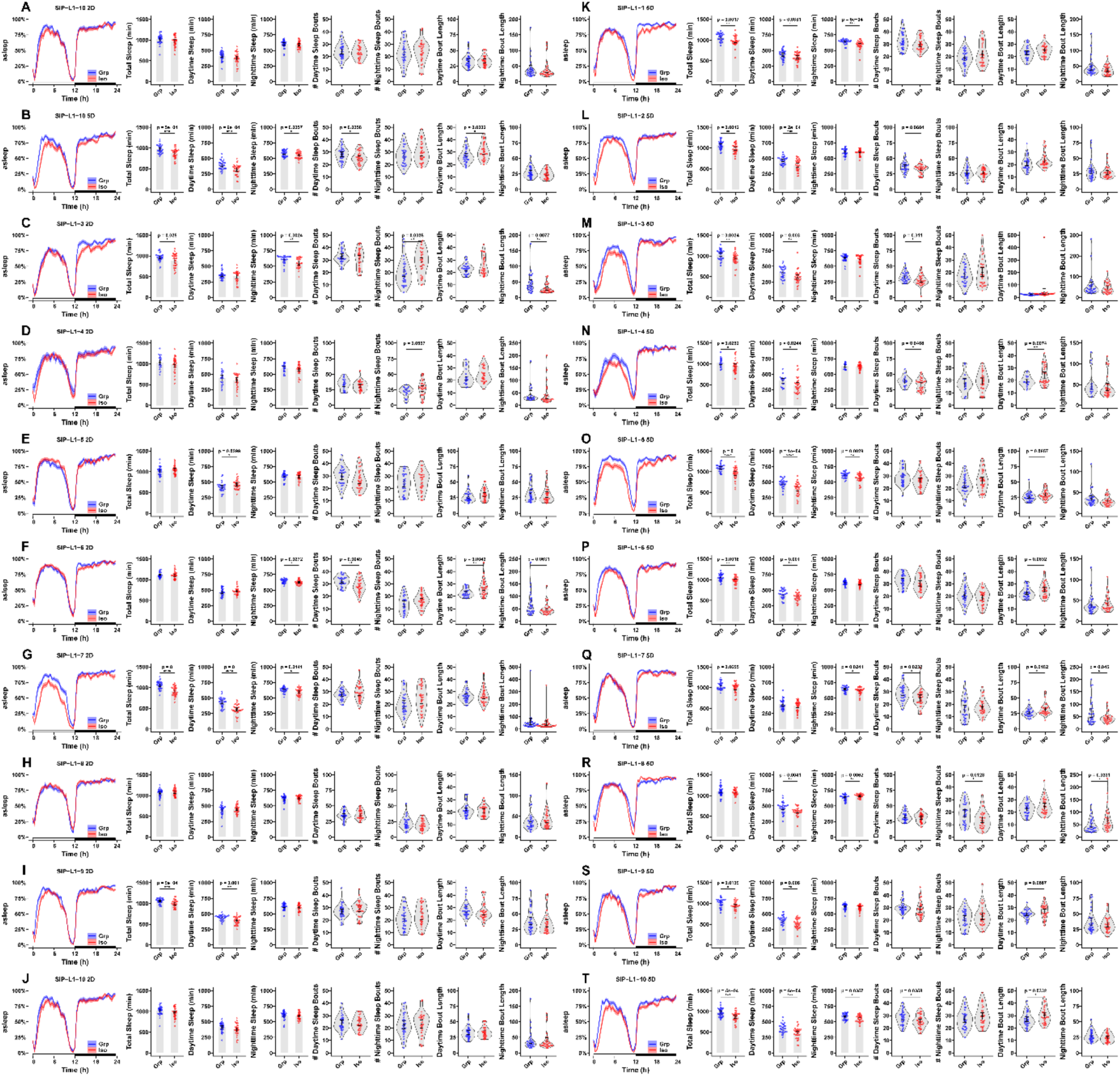
Sleep profiles and parameters for SIP-L1 Strains. **(A–T)** SIP-L1 strains 1–10’s sleep profiles, sleep parameters, and bout distributions after 2 days **(A–J)** or 5 days **(K–T)** of treatment. Statistical significance was assessed by two-sided unpaired *t*-tests (*\*P<0.05, **P<0.01, ***P<0.001, ****P<0.0001*; *n* = 17–32).

**Supplementary Figure 4.**
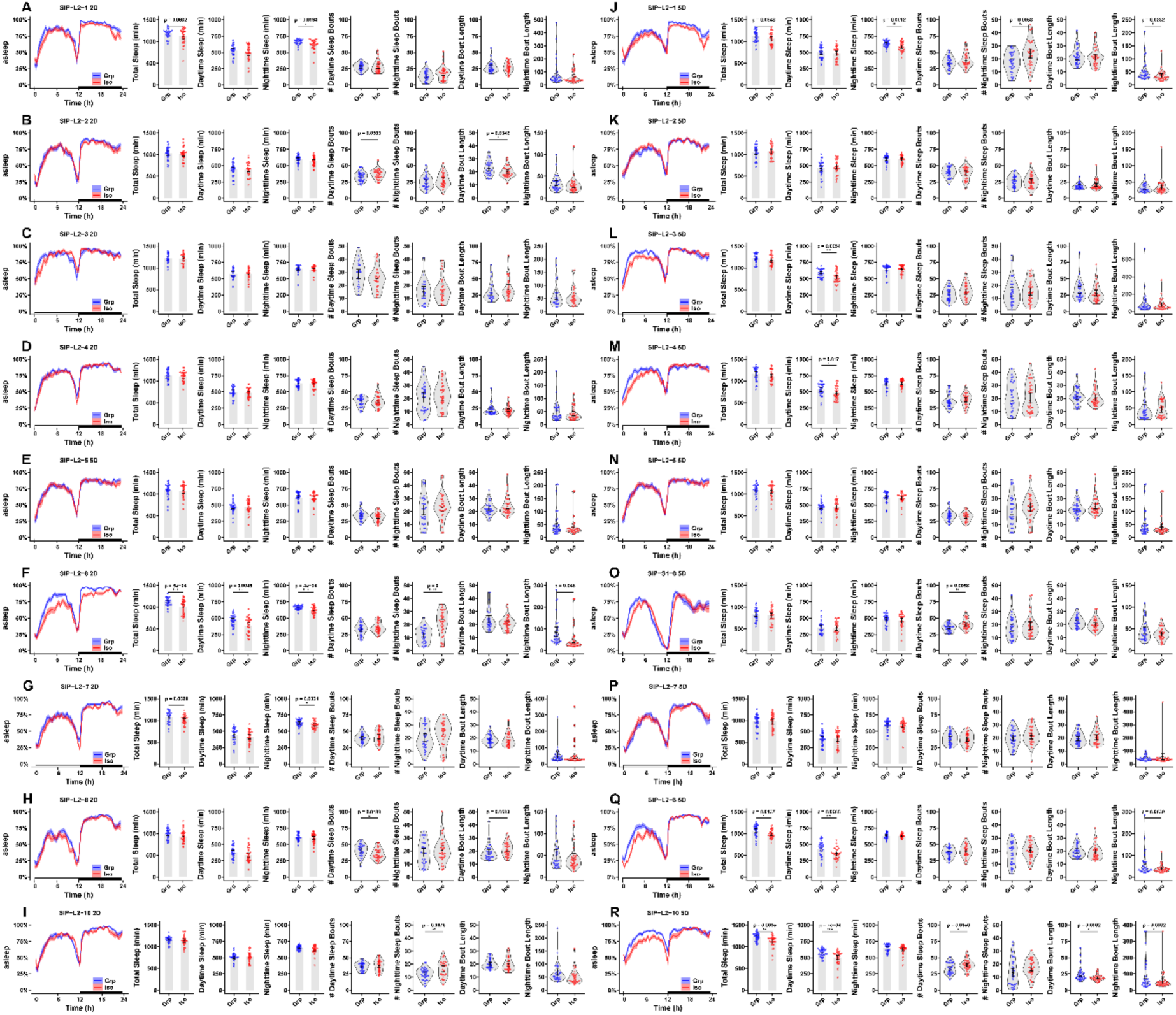
Sleep profiles and parameters for SIP-L2 Strains. **(A–R)** SIP-L2 strains 1–8, and 10’s sleep profiles, sleep parameters, and bout distributions after 2 days **(A–I)** or 5 days **(J–R)** of treatment as described. Statistical significance was assessed by two-sided unpaired *t*-tests (*\*P<0.05, **P<0.01, ***P<0.001, ****P<0.0001*; *n* = 25–32).

**Supplementary Figure 5.**
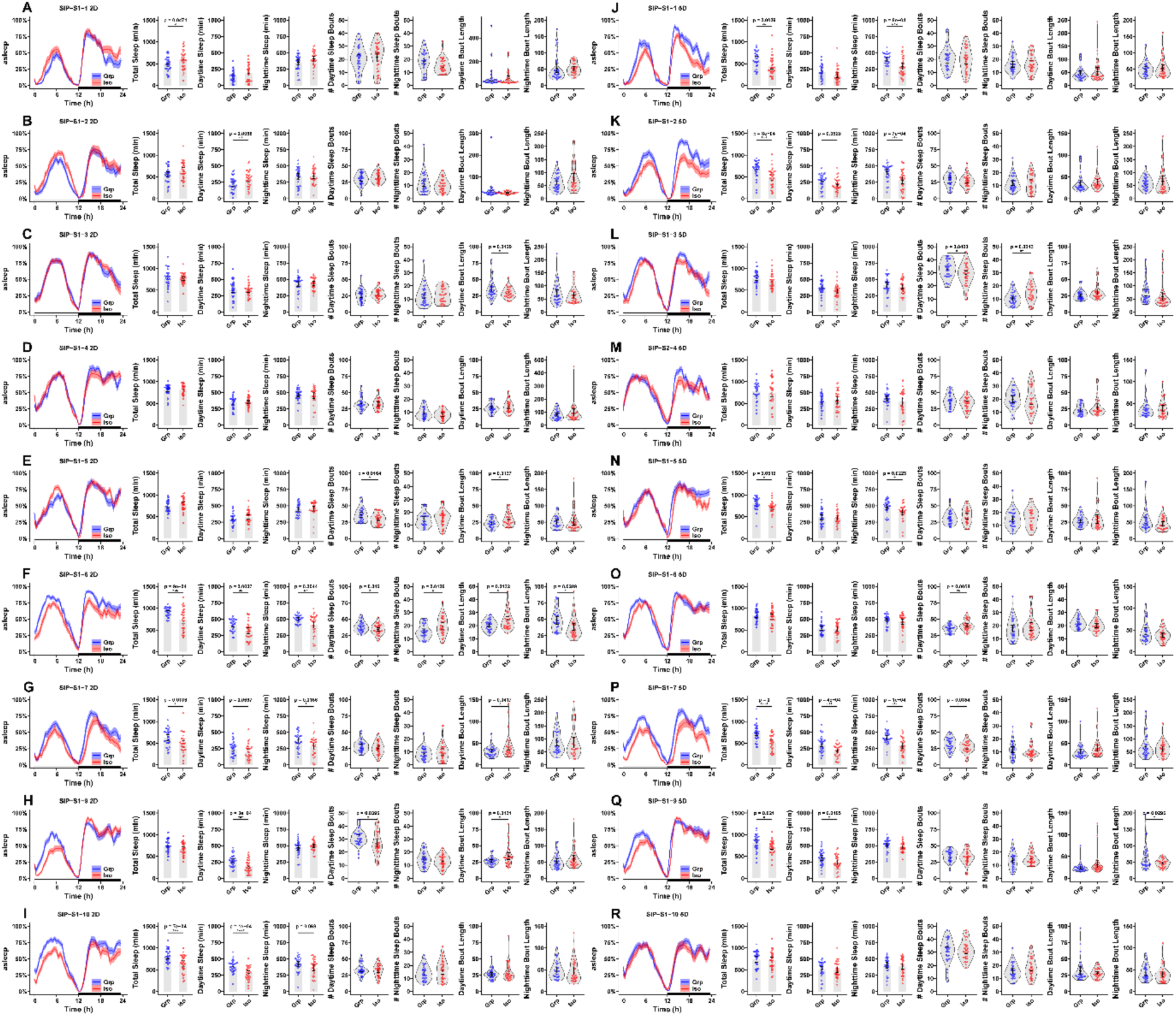
Sleep profiles and parameters for SIP-S1 Strains. **(A–R)** SIP-S1 strains 1–7, and 9–10’s sleep profiles, sleep parameters, and bout distributions after 2 days **(A–I)** or 5 days **(J–R)** of treatment. Statistical significance was assessed by two-sided unpaired *t*-tests (*\*P<0.05, **P<0.01, ***P<0.001, ****P<0.0001*; *n* = 26–32).

**Supplementary Figure 6.**
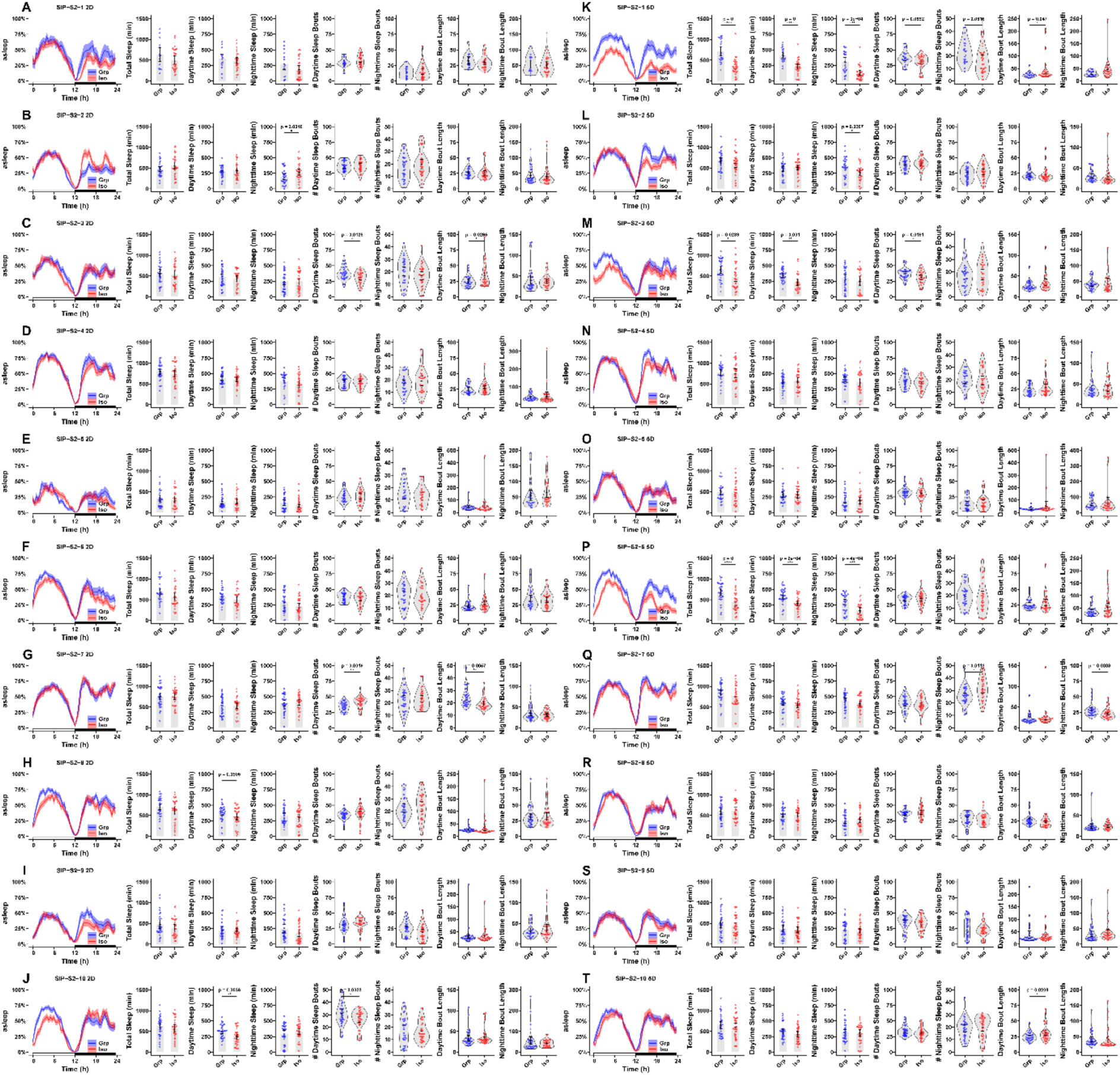
Sleep profiles and parameters for SIP-S2 Strains. **(A–T)** SIP-S2 strains 1–10’s sleep profiles, sleep parameters, and bout distributions after 2 days **(A–J)** or 5 days **(K–T)** of treatment. Statistical significance was assessed by two-sided unpaired *t*-tests (*\*P<0.05, **P<0.01, ***P<0.001, ****P<0.0001*; *n* = 15–32).

## References

1. Cacioppo, J.T., et al., Do lonely days invade the nights? Potential social modulation of sleep efficiency. Psychol Sci, 2002. 13(4): p. 384–7.

2. Holt-Lunstad, J., Social Connection as a Public Health Issue: The Evidence and a Systemic Framework for Prioritizing the “Social” in Social Determinants of Health. Annu Rev Public Health, 2022. 43: p. 193–213.

3. Kurina, L.M., et al., Loneliness is associated with sleep fragmentation in a communal society. Sleep, 2011. 34(11): p. 1519–26.

4. Vora, A., et al., The impact of social isolation on health and behavior in Drosophila melanogaster and beyond. Brain Science Advances, 2022. 8(3): p. 183–196.

5. Li, W., et al., Chronic social isolation signals starvation and reduces sleep in Drosophila. Nature, 2021. 597(7875): p. 239–244.

6. Gil-Martí, B., et al., Socialization causes long-lasting behavioral changes. Sci Rep, 2024. 14(1): p. 22302.

7. Ganguly-Fitzgerald, I., J. Donlea, and P.J. Shaw, Waking experience affects sleep need in Drosophila. Science, 2006. 313(5794): p. 1775–81.

8. Serrano Negron, Y.L., N.F. Hansen, and S.T. Harbison, The Sleep Inbred Panel, a Collection of Inbred Drosophila melanogaster with Extreme Long and Short Sleep Duration. G3 Genes|Genomes|Genetics, 2018. 8(9): p. 2865–2873.

9. Harbison, S.T., et al., Selection for long and short sleep duration in Drosophila melanogaster reveals the complex genetic network underlying natural variation in sleep. PLOS Genetics, 2017. 13(12): p. e1007098.

10. Ghosh, A. and V. Sheeba, VANESSA-Shiny Apps for Accelerated Time-series Analysis and Visualization of Drosophila Circadian Rhythm and Sleep Data. J Biol Rhythms, 2022. 37(2): p. 222–231.

11. Persons, J.L., et al., PHASE: An Open-Source Program for the Analysis of DrosophilaPhase, Activity, and Sleep Under Entrainment. J Biol Rhythms, 2022. 37(4): p. 455–467.

12. Geissmann, Q., et al., Rethomics: An R framework to analyse high-throughput behavioural data. PLOS ONE, 2019. 14(1): p. e0209331.

13. Sisobhan, S., et al., SleepMat: a new behavioral analysis software program for sleep and circadian rhythms. Sleep, 2022. 45(12): p. zsac195.

14. Vecsey, C.G., et al., Analysis of Sleep and Circadian Rhythms from Drosophila Activity-Monitoring Data Using SCAMP. Cold Spring Harb Protoc, 2024. 2024(11): p. pdb.prot108182.

15. Silva, R.F.O., et al., Automated analysis of activity, sleep, and rhythmic behaviour in various animal species with the Rtivity software. Scientific Reports, 2022. 12(1): p. 4179.

16. Gilestro, G.F. and C. Cirelli, pySolo: a complete suite for sleep analysis in Drosophila. Bioinformatics, 2009. 25(11): p. 1466–7.

17. Wickham, H., et al., dplyr: A Grammar of Data Manipulation. 2023.

18. Wickham, H., ggplot2: Elegant Graphics for Data Analysis. 2016: Springer-Verlag New York.

19. Clarke, E., S. Sherrill-Mix, and C. Dawson, ggbeeswarm: Categorical Scatter (Violin Point) Plots. 2023.

20. Dawson, C., ggprism: A ’ggplot2’ Extension Inspired by ’GraphPad Prism’. 2024.

21. Kassambara, A., ggcorrplot: Visualization of a Correlation Matrix using ’ggplot2’. 2022.

